# Timing seed germination under changing salinity: a key role of the ERECTA receptor-kinases

**DOI:** 10.1101/576512

**Authors:** Amrit K. Nanda, Abdeljalil El Habti, Charles Hocart, Josette Masle

## Abstract

Appropriate timing of seed germination is crucial for the survival and propagation of plants, and for crop yield, especially in environments prone to salinity or drought. Yet, how exactly seeds perceive changes in soil conditions and integrate them to trigger germination remains elusive, especially once non-dormant. Here we report that the *Arabidopsis* ERECTA (ER), ERECTA-LIKE1 (ERL1) and ERECTA-LIKE2 (ERL2) leucine-rich-repeat receptor-like kinases synergistically regulate germination and its sensitivity to salinity and osmotic stress. Loss of *ER* alone, or in combination with *ERL1* and/or *ERL2* slows down the initiation of germination and its progression to completion, or arrests it altogether until better conditions return. That function is maternally controlled via the embryo surrounding tissues, primarily the properties of the seed coat determined during seed development on the mother plant, that relate to both seed coat expansion and subsequent differentiation, particularly the formation of its mucilage. Salt-hypersensitive *er, er erl1, er erl2* and triple mutant seeds also exhibit increased sensitivity to ABA during germination, and under salinity show an enhanced upregulation of the germination repressors and inducers of dormancy *ABA-insensitive-3, ABA-insensitive-5*, DELLA encoding *RGL2* and *Delay-Of-Germination-1*. These findings reveal a novel role of the ERECTA kinases in the sensing of conditions at the seed surface and the integration of developmental and stress signalling pathways in seeds. They also open novel avenues for the genetic improvement of plant adaptation to harsh soils.

**Highlight:** The ERECTA family of receptor-kinases regulates seed germination under salinity, through mucilage-mediated sensing of conditions at the seed surface, and interaction with secondary dormancy mechanisms.

## Introduction

Seed germination is a vital life-cycle transition in plants. When and under which conditions it occurs, largely determine survival, reproductive success, yield and ability to expand. To maximise chances of timely germination under favourable conditions, seeds have evolved mechanisms for dormancy, a state that prescribes the environmental conditions that need to occur before germination can take place (Bewley, 1997; Baskin and Baskin, 2004; Finch-Savage and Leubner-Metzger, 2006). As dormancy fades over time, or is lifted by appropriate cues, water becomes the most critical requirement for successful germination, in interaction with temperature (Alvarado and Bradford, 2002). As soon as moisture contacts it, the highly desiccated seed imbibes like a sponge. Seed imbibition reactivates metabolism and enables embryo expansion and rupture of its protective tissues. In *Arabidopsis thaliana*, this occurs in two separate steps, the rupture of the testa, or seed coat, a dead tissue, and then endosperm rupture (Liu et al., 2005; Müller et al., 2006). Embryo reactivation and weakening of surrounding tissues are tightly coordinated, through complex biochemical and hormonal pathways, with a prominent role of abscisic acid (ABA) and gibberellins (GAs), in interaction with ethylene, brassinosteroids and reactive oxygen species (ROS) (Koornneef and Van der Veen, 1980; Steber and McCourt, 2001; Bailly, 2004; Finch-Savage and Leubner-Metzger, 2006; Kucera et al. 2005; Finkelstein et al., 2008; Weitbrecht et al., 2011; Rajjou et al., 2012; Yu et al., 2016). ABA inhibits germination whereas GAs promote it, through regulation of inter-signalling between seed coat, endosperm, and embryo, in a feedback loop involving DELLA proteins and interactions with cell-wall remodelling enzymes (Müller et al., 2006; Stamm et al., 2012; Graeber et al., 2014; Nonogaki, 2014).

Drought and salinity stress are two inter-related and widespread conditions in natural environments, and major causes of germination failure, poor crop establishment and yield loss (Boyer, 1982; Bradford K.J., 1990; Finch-Savage and Leubner-Metzger, 2006; Yamaguchi and Blumwald, 2005; Munns and Tester, 2008). The high vulnerability of seeds to these stresses has long-been recognised. Yet, the molecular controls remain poorly understood, apart from evidence for a deregulation of ABA-GA homeostasis and an impairment of ethylene and ROS signalling (Lopez-Molina et al., 2001; Kim et al., 2008; Yuan et al. 2010; Yu et al., 2016). Natural genetic variation in seed germination under optimal conditions, drought or salinity has been widely documented, and numerous QTLs identified (e.g. Quesada et al., 2002; Clerkx et al., 2004; Galpaz and Reymond, 2010; Wang et al., 2010; DeRose-Wilson and Gaut, 2011; Yuan et al., 2016). This demonstrates the potential for genetic improvement, but also the complexity of the underlying molecular pathways. While the genetic dissection of seed dormancy has received much attention, very few genes have been demonstrated to control the germination of non-dormant seeds to tune it to prevailing soil conditions (Kim et al., 2008; Ren et al., 2010; Yu et al., 2016). How seeds monitor their surroundings, how this information is communicated to their inner compartments and modulates the intricate communication between them and the environment that timely germination requires, remains little known (Donohue et al., 2010).

Receptor-like protein kinases (RLKs) at the cell plasma membrane play major roles in signal perception and transduction to downstream intra- and inter-cellular signalling networks. A vast array of RLKs are encoded by plant genomes (Shiu et al., 2004). Among them are Leucine-Rich-Repeat Receptor-Like Kinases (LRR-RLKs) which form a large family of receptor proteins characterised by an extra-cellular receptor domain, a trans-membrane domain and an intra-cellular kinase domain for signal transduction through phosphorylation cascades. The few that have been characterised provide evidence for central functions in integrating developmental, hormonal, and abiotic stress or defence signalling pathways (Becraft, 2002; Osakabe et al., 2013). Scant information is available on RLKs in seeds, even though developing seeds show high abundance of secreted peptides and recent studies point to the importance of peptide-mediated signalling in inter-compartmental coordination during seed development (Ingram and Gutierrez-Marcos, 2015).

The *Arabidopsis ERECTA* gene family (*ER*f) encodes three closely related LRR-RLKs - *ER, ERL1* and *ERL2*-known to synergistically regulate many aspects of plant development and morphogenesis with prominent roles in organ shape, stomatal patterning, cell proliferation and meristematic activity (Torii et al., 1996; Shpak et al., 2004; 2005; Pillitteri et al., 2007; Uchida et al., 2012; Bemis et al., 2013; Etchells et al., 2013; Ikematsu et al., 2016), as well as being involved in some pathogenic responses (Godiard et al., 2003; Llorente et al., 2005; Jordá et al., 2016). In contrast, little is known of its function in abiotic stress responses, beyond a role in leaf heat tolerance (Shen et al. 2015). We earlier reported a role of ERECTA as a major controller of water use efficiency, under both well watered and drought conditions (Masle et al., 2005). That function appears to be broadly conserved in diverse species (Xing et al., 2011; Zheng et al., 2015), and is suggestive of an important adaptive role of the ERf to abiotic stress. Here we probe the ERf function during germination, a key switch that is extremely sensitive to variations in osmotic and ionic soil conditions, both of which vary widely in nature.

## Material and Methods

### Plant material and growth conditions

*Arabidopsis thaliana* Columbia (Col-0, CS1093) was used as wild type (WT), alongside two independent sets of single, double and whole ERECTA family loss-of-function mutants: one carrying the previously characterised mutations *er105, erl1-2, erl2-1*, in the *ER* (At2g26330), *ERL1* (At5g62230) and *ERL2* (At5g07180) genes, respectively (Torii et al., 1996; Shpak et al., 2004; Masle et al., 2005; Bundy et al., 2012; Bemis et al., 2013) and here coded *er, erl1* and *erl2* for simplicity; the other carrying the *er2* (*C3401*) mutation (Rédei, 1992; Lease et al., 2001; Masle et al., 2005; Hall et al., 2007), and the *erl1-5* (SALK_019567) and *erl2-2* (SALK_015275C) insertional mutations from the SALK Institute collection (Alonso et al.2007). Absence of residual target gene expression in the latter two lines was confirmed, and T-DNA insertion sites verified (insertion located 4605 bp and 2775 bp from *ERL1* and *ERL*2 start codon in *erl1-5* and *erl2-2*, respectively).

Double and triple *er*f mutants were generated through crosses. As the triple mutants are sterile, the segregating progeny of *er erl1/+ erl2*, or *er2 erl1-5+/-erl2-2* was used to investigate germination of triple mutant seeds, and is referred to in text and figures as *er erl1/seg erl2* or *er2 erl1-5/seg erl2-2*, respectively.

All seeds in any given experiment were of the same age and harvested from spaced plants grown together, under the same conditions (21°C constant temperature; 12 or 16 h day length, depending on experiment; 120-130 μmol quanta m^−2^ s^−1^ light intensity). For investigation of parent-of-origin effects on seed germination and seed size, seeds were manually excised from tagged mature siliques, of the same age and same position on the primary inflorescence.

### Germination assays

All assays were done using seeds stratified by moist chilling at 4°C to remove residual dormancy. Seeds were surface-sterilised and sown on 0.7% agar media supplemented with Hoagland’s nutrient solution (2 mM KNO_3_, 5 mM Ca[NO_3_]_2_4H_2_O, 2 mM MgSO_4_7H_2_O, 2 mM KH_2_PO_4_, 0.09 mM Fe-EDTA and micronutrients) pH 5.8, and NaCl or KCl in desired concentrations. For germination assays under iso-osmotic conditions generated by PEG8000 or NaCl, seeds were plated on filter paper imbibed with solutions of NaCl or PEG8000 dissolved in water. The osmotic pressure (π_e_) of the basal medium or NaCl- or KCl-containing media was calculated using the classic van‘t Hoff equation and verified experimentally using a VAPRO vapour pressure osmometer (Wescor Inc.). The concentrations of PEG8000 required to obtain a given π_e_ were determined from a calibration curve of π_e_ as a function of [PEG] using the same instrument. Seeds WT and all *er*f mutant combinations were sown in equal number (n≥33) within each of 3 to 4 plates (total n = 100 to 120 seeds per line per treatment and experiment). After stratification at 4°C, in the dark for 2-3 days, plates were exposed to continuous light (100-115 μmol quanta m^−2^ s^−1^) and a constant 21°C temperature. “Demucilaged” seeds were sown straight after mucilage removal (see protocol below), and kept at 4°C in darkness for an additional day, so as to keep total stratification time to 48 h, as for control intact seeds.

Seeds were individually scored for both testa and endosperm rupture (germination *sensu stricto*) under a binocular microscope, within the growth chamber, and at 3-4 h intervals until all seeds on control plates (0 mM NaCl) had germinated (i.e. 30 hours at most), or three times to once daily, as appropriate on NaCl, KCl, or PEG plates, until no change in scores was observed. Data are represented either as percentages of seeds exhibiting testa or endosperm rupture as a function of incubation time post-stratification, or as T_50_ values, corresponding to the times (h post-stratification) when 50% of seeds showed testa or endosperm rupture (Bewley et al., 2013).

### Embryo culture

Mature embryos were excised from dry seeds pre-imbibed with water for 1-2 h, briefly rinsed twice in water to remove endosperm debris and plated on either 0 or 150 mM NaCl media, placed in the dark at 4°C for 3 d, before transfer to the growth chamber. Embryos were individually imaged at the time of transfer and again 72h later using a LEICA M205 FA microscope fitted with a DFC 550 camera (LEICA Instruments). Relative embryo expansion rates over that 72 h interval were calculated from measurements of projected areas using *Image*J software.

### Staining procedures

GUS histochemical staining of seeds from *proERf:GUS* reporter lines (Shpak et al., 2004) was performed on embryos dissected from dry and germinating seeds sampled from 0 and 150 mM NaCl plates. Staining was done as described (Sessions et al., 1999).

For tetrazolium permeability assays (Debeaujon and Koornneef, 2000) dry seeds were incubated in the dark in an aqueous solution of 1% (w/v) tetrazolium red (2,3,5-triphenyltetrazolium chloride, Sigma-Aldrich) at 30°C for 4, 24, 48, 72 and 120 h, and then rinsed twice with deionised water, resuspended in 95% ethanol and quickly ground to extract formazans. The final volume was adjusted to 2 ml with 95% ethanol, followed by centrifugation at 15000 *g* and measurement of supernatant’s absorbance at 485 nm, using a Tecan Infinite M1000 Pro spectrophotometer (Tecan Trading AG, 2008). Each sample was assayed in triplicates.

Mucilage ruthenium red staining was performed as described (http://www.bio-protocol.org/e1096). Ruthenium red stains acidic pectins (Hanke and Northcote, 1975) and is widely used to stain *Arabidopsis* seed mucilage (Western et al., 2000; Penfield et al., 2001).

### Profiling fatty acid methyl esters derived from lipids stored in the embryo

Fifty mature embryos were dissected from dry seeds after 1 h imbibition in water, in 4 replicates per genotype. Fatty acid methyl esters (FAMEs) were prepared by direct transesterification as described by James et al. (2011). Embryos were placed in a reacti-vial (1.5mL) fitted with a Teflon-lined cap. To this was added CHCl_3_ (50 µL) followed by the internal standard, heptadecanoic acid (C17:0, 15 µL, 9.66 mg in 25mL CHCl_3_), and methanolic HCl (3M, 500 µL). The samples were mixed and heated at 90 °C for 60 min, and then allowed to cool before being washed into glass tubes with CHCl_3_. Water (1 mL) was added to each tube and the FAMEs extracted (hexane:chloroform, 4:1 v/v, 3 x 1 mL). The extracts were combined and washed with water (200 µL). The organic phase was then dried with anhydrous Na_2_SO_4_, decanted and evaporated under nitrogen. The residue was dissolved in CH_2_Cl_2_ (150 µL) and transferred to GC/MS auto-sampler vials for analysis.

### Mucilage extraction and analysis

Mucilage extraction was performed on aliquots of 40 mg dry seeds. Each aliquot (n=4 per genotype per experiment) was suspended in 1ml milliQ water, followed by shaking at 500 rpm for 24 h at 4°C, vortex for 5 s, and centrifugation at 8000 *g* for 3 mins. 600 µl supernatant was recovered. Seeds were rinsed twice with 200 µl water, and 200 µl supernatant was recovered after vortexing and centrifugation. The pooled supernatants (1ml total volume) was snap-frozen in liquid nitrogen and immediately lyophilised. The mucilage thus recovered was weighed on a 10^−6^ g high precision micro-balance. Given the observed genetic variation in seed size (see Text), sub-aliquots of a known number of seeds (at least 500) were weighed, imaged at high resolution and analysed for size with *Image*J prior to mucilage extraction, allowing derivation of average mucilage amount per seed. The reductions of uronic acid methyl-esters and free uronic acids in the extracted mucilage were carried out following established protocols (Kim and Carpita, 1992; Pettolino et al., 2012). The reduced polysaccharides were then hydrolysed, reduced, acetylated and subjected to GC/MS analysis as described (Peng et al., 2000).

### Analysis of seed sodium content

Dry seeds (3 biological replicates of 10 mg seeds each were per genotype and treatment) were imbibed and stratified at 4°C in the dark in a 0 or 150 mM NaCl solution for two days followed by 24 h at room temperature with shaking. Seeds were rinsed 3 times with 2 ml water, freeze-dried, weighed and microwave-digested for 2 h in 4 ml of 20% nitric acid at 175°C (USAP Method 3051). Digest volumes were diluted to a final volume of 5 ml. Sodium ions were measured by ICP-OES (Varian Vista-Pro CCD Simultaneous).

### Quantitative RT-PCR

Total RNA was extracted from dry, imbibed or germinating seeds using TRIzol reagent (Invitrogen). mRNA isolation and reverse transcription were done as described (Chen et al., 2018). Primer sequences are given in Supplementary Table 1. The analysis was done on four biological replicates per genotype, time point and treatment, of 300 seeds each, sampled from 4 plates where all genotypes compared were represented. Target gene expression levels were normalised to the geometric mean of expression levels of four reference genes, *APT1* (At1g27450), *PDF2* (At1g13320), *bHLH* (At4g38070), and *PPR* (At5g55840). Gene expression was measured just before sowing (“Dry” seeds) and then at: the end of seed imbibition and stratification (germination stage I); 20 h later (stage II, testa rupture); and 52 h later (stage III-G, endosperm rupture; seeds non-germinated yet, III-NG, were analysed separately). Seeds were sampled within the cold room or growth room (dry seeds and stage I to III, respectively), within 5 minutes from start to finish for each plate, and immediately snap-frozen in liquid nitrogen. The experiment was repeated three times.

### Statistical analysis

Statistical significance of results was analysed using the Statistix 9 software (Analytical Software, Tallahassee, USA). For multivariate comparison of mucilage composition profiles, discriminant Orthogonal Projected Latent Structure (OPLS) analysis was carried out using the SIMCA software (Umetrics, www.umetrics.com) with salinity as a quantitative variable.

## Results

### The ERf controls the timing and pace of germination in response to changing salinity and osmotic conditions

Loss of ER/ERL function had no effect on testa nor endosperm rupture on 0 mM NaCl media, except in *er erl1* seeds which showed a small but systematic lag in testa rupture (Fig. 1 and Supplementary Fig. S1A-B). That lag carried through to the next germination phase leading to radicle protrusion. Salinity delayed germination in a dose-dependent manner (Supplementary Fig. S2A, B), as expected, but with striking differences among lines (Fig. 1 and Supplementary Fig. S1C-D). Wild type (WT), *erl1, erl2* and *erl1 erl2* seeds germinated first, ahead of *er, er erl2, er erl1* and finally *er erl1/seg erl2* seeds, due to both delayed testa rupture and slower progression to endosperm rupture. Similar results were obtained with an independent set of *er*f knock-out mutants carrying different *er*f null alleles (Supplementary Fig. S3). This demonstrates that the observed genetic variations in seed germination are causally related to disruption of the *ER*f genes. Similar germination kinetics were also obtained regardless of whether seeds were challenged with salinity stress post-stratification or directly from sowing (Fig. 2A-D). Strikingly, when exposed to 150 mM NaCl once germinated, all genotypes displayed similar sensitivity to salinity stress (Fig. 2E). Together, these data demonstrate a germination-specific function of the ERf in the sensing and signaling of salinity stress. That function requires ER but involves the three family members, in a non-totally redundant manner.

**Figure 1.**
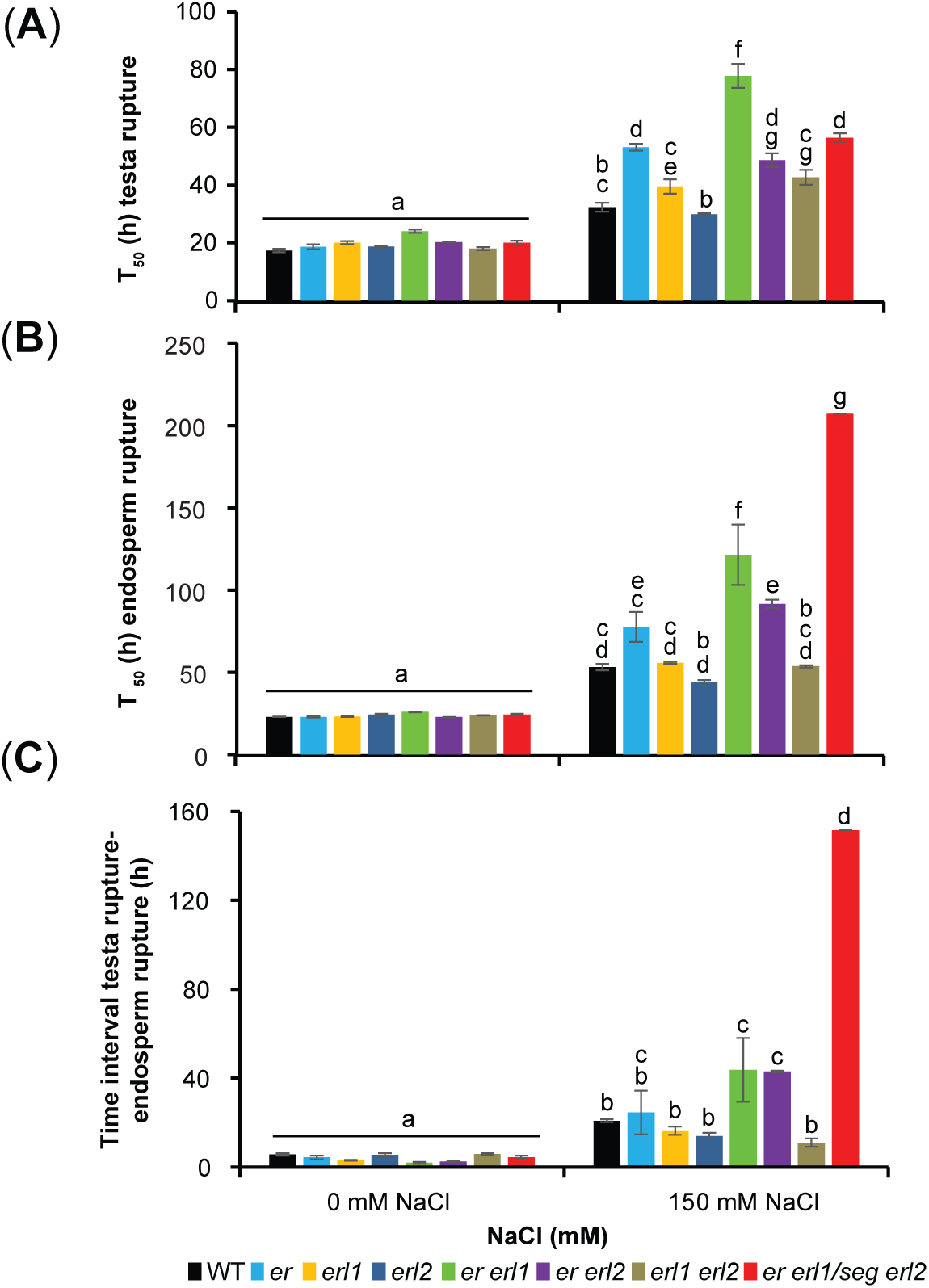
The three ERECTA family members synergistically control the timing and pace of seed germination under salinity. **A, B**, T_50_ values (h post-stratification) for testa rupture (**A**) and endosperm rupture (**B**). **C**, Time interval (h) between the two steps. Experiment repeated 5 times with different seed batches. As the triple mutant is sterile, the segregating progeny of *er erl1+/-erl2* plants was used to investigate germination of triple mutant seeds, and is referred to in text and figures as *er erl1/seg erl2*. **A-C**, Values are means and s.e.m. (n = 4 plates, 30 seeds per genotype per plate). Different letters above bars denote significant differences by two-way ANOVA and Tukey HSD pair-wise tests (*P*□< □0.001).

**Figure 2.**
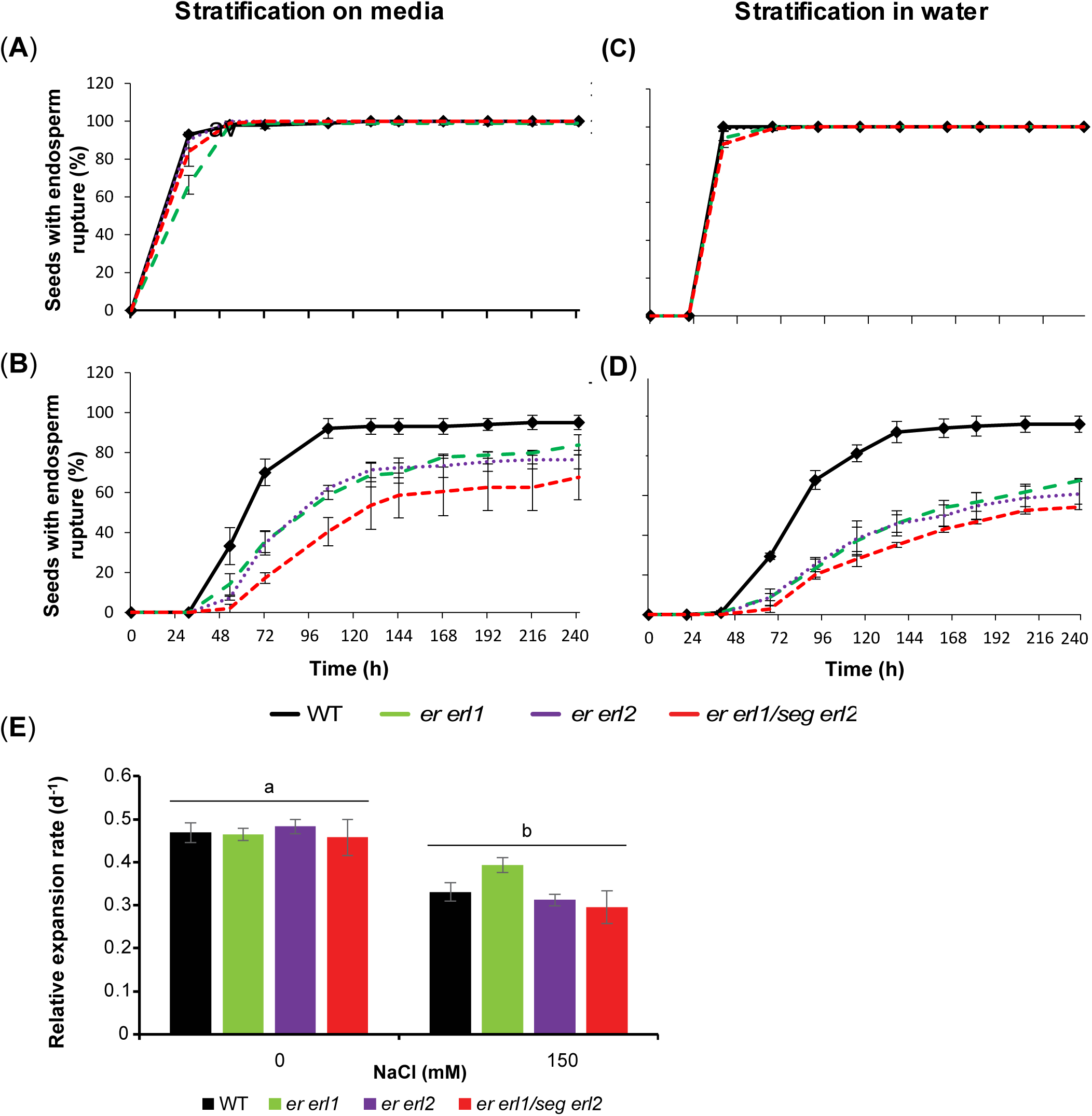
Germination-specific function of the ERf on sensitivity to salinity. **A-D**, Time-course of endosperm rupture for *er, er erl1, er erl2* or *er erl1/seg erl2* seeds over a 10 d incubation period on 0 **(A, C)** or 150 mM NaCl agar media **(B, D)** following imbibition and stratification either directly on media **(A, B)** or in water prior to plating **(C, D). E**, Seedling relative expansion rates (d^-1^) on 0 or 150 mM NaCl media. Seeds were first germinated on NaCl-free media and then transferred to fresh 0 mM or 150 mM NaCl plates for monitoring their expansion over the next 72 h, measurements of whole seedling projected area on images captured using *ImageJ*. Different letters indicate significant differences by two-way ANOVA and Tukey HSD pair-wise tests (*P*□<□0.001), n = 7.

*ER*f expression during seed germination has not been reported. To investigate it, we examined *ER*f promoter activity in transgenic seeds expressing *proERf:GUS* constructs (Supplementary Fig. S4). *ER*f expression patterns did not appear to be influenced by salinity, but differed among family members, with *ERL2* expression seen only in the cotyledons and the shoot apical meristem, while *ER* and *ERL1* promoter activities were also detected in the hypocotyl. Measurements of transcript abundance by RT-qPCR (Fig. 3) confirmed the presence of *ER*f transcripts in dry seeds and showed a strong and early induction of *ER* and *ERL1* expression during stratification and imbibition (germination phase I), and the next phase (stage II) leading to testa rupture, while *ERL2* remained lowly expressed. Salinity induced *ER* expression, especially during germination phase III, leading to radicle protrusion, but had little influence on *ERL1* or *ERL2* expression. These results support a role of the ERf throughout germination, with specificity among family members.

**Figure 3.**
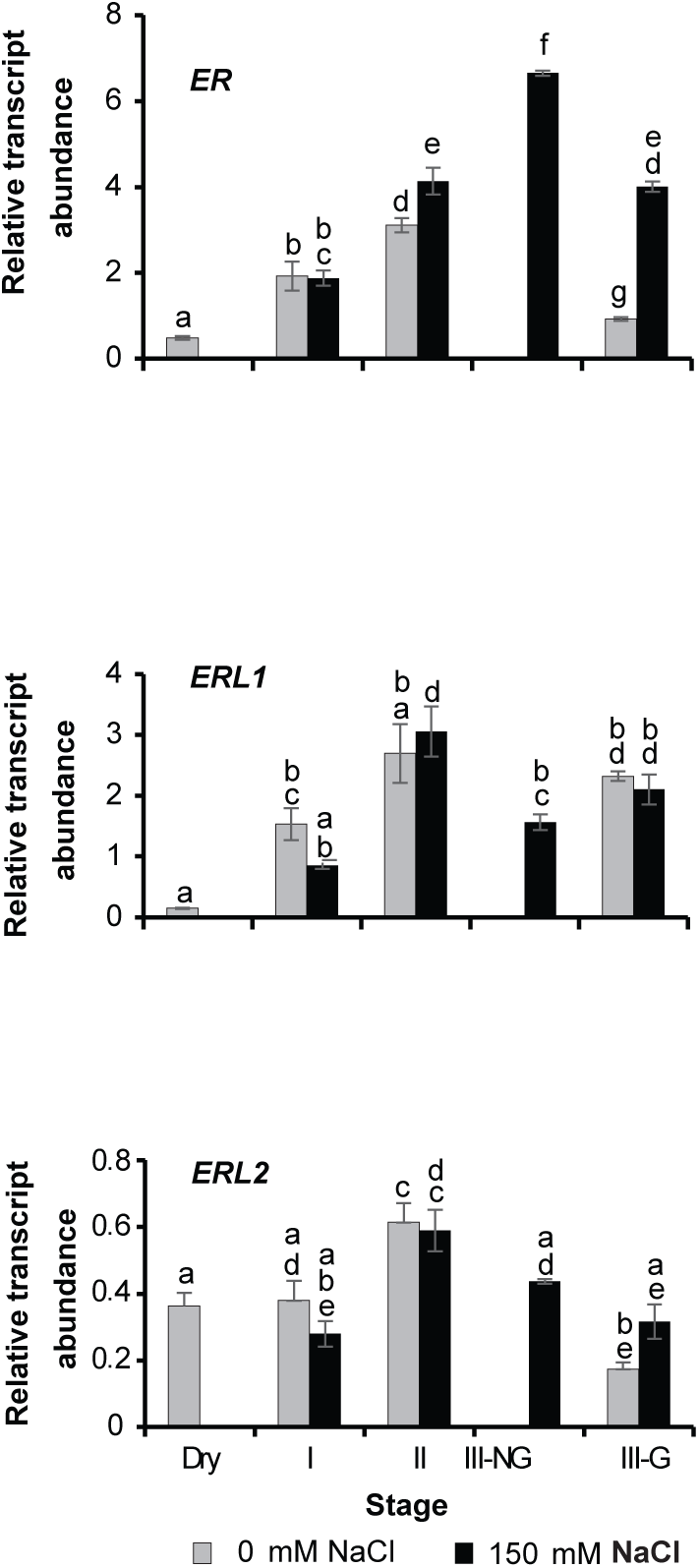
*ER*f transcripts are present in mature dry seeds and *ER*f *de novo* transcription is activated early during germination. *ER*f gene expression in WT dry seeds (“Dry”) and germinating seeds at: the end of imbibition and stratification (stage I); 20 h later (stage II, testa rupture); and 52 h later (stage III-G, endosperm rupture completed on control media). Non-germinated seeds at that time on 150 mM NaCl (label III-NG) were sampled and analysed separately. Different letters indicate significant differences by two-way ANOVA and Tukey HSD pair-wise tests (*P*□< □0.001), n=4 seed pools per genotype and treatment, of 300 seeds each. The experiment was repeated 3 times.

Salinity induces both osmotic and ionic stress (Munns and Tester, 2008). To investigate the contributions of these two components, we next examined germination responses to Polyethylene Glycol (PEG)8000 - a high molecular weight non-permeating osmoticum mimicking drought-induced osmotic stress-, in the salinity hyper-sensitive mutants, *er, er erl1, er erl2* and *er erl1/seg erl2*. Under iso-osmotic external conditions (media osmotic pressure, π_e_), seed germination was significantly less inhibited by PEG than NaCl. Up to 0.50 MPa π_e_ (equivalent to 100 mM NaCl), PEG was innocuous (Fig. 4A). When, however, PEG was provided at higher concentrations raising π_e_ to 0.74 and 0.99 MPa (iso-osmotic conditions with 150 and 200 mM NaCl, respectively), germination was slowed down but to a greater extent in the double and triple mutants than WT. Nevertheless, the germination delay was mild, of the order of 1 day. By d3 post-stratification, germination was complete (WT, *er, er erl1* and *er erl2* seeds) or near complete (*er erl1/seg erl2* seeds, 90% germination), even under 0.99 MPa (Fig. 4A-B), in contrast to the strong to total germination inhibition observed under 200 mM Nacl, at the same π_e_ (Fig. 4C, Supplementary Fig. S2A,B). Only at much higher PEG concentrations was as severe an inhibition observed, but some seeds still germinated (Fig. 4B). Taken together, these data indicate that 1) in the germination-permissive range of NaCl concentrations, the ERf modulates seed germination sensitivity to salinity mostly via interactions with NaCl ionic effects; 2) however, the ER is also involved in the control of germination sensitivity to osmotic and hyper-osmotic stress.

**Figure 4.**
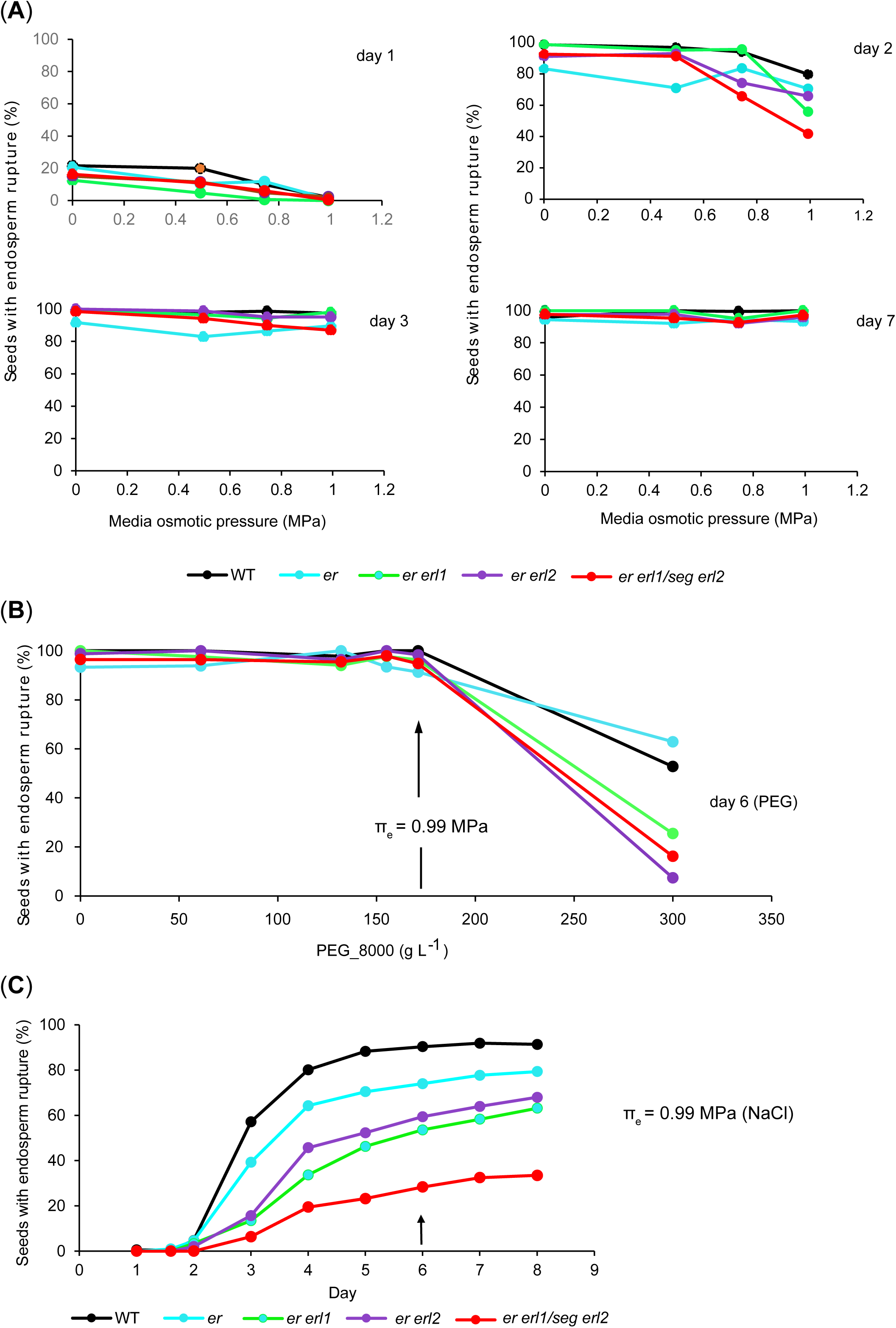
The ERf regulates seed germination sensitivity to salinity mostly via interactions with its ionic effects, but is also involved in the control of germination under osmotic stress. **A**, Percentage of seeds with endosperm rupture for WT and the NaCl hyper-sensitive mutants *er erl1, er erl2*, and *er erl1/seg erl2*, on day 1, 2, 3 and 7 post-stratification as a function of media osmotic pressure (π_e_) varied through supplementation of PEG_8000 at concentrations ranging from 0 to 171g L^-1^. n=100-200 seeds per replicate. **B**, Germination response over an extended range of PEG concentrations, in an independent experiment with a different seed batch. Data points depict the percentage of seeds exhibiting endosperm rupture 6d post-stratification. n= 100-200 seeds per replicate. **C**, Kinetics of seed germination under 0.99 MPa π_e_ induced by NaCl. Same seed batch as in (**B)**. The arrow points to germination scores on d6 when, under iso-osmotic conditions induced by PEG, at least 90% seeds had germinated (see panel **B**). n = 3 plates, 30 seeds per plate and per genotype. Experiments replicated 3 times.

The NaCl-hypersensitive *er*f mutants also exhibited increased sensitivity to KCl, but to a much lower extent than to NaCl under iso-osmotic conditions (Fig. 5). This result indicates that the ERf function in seed germination under salinity predominantly relates to effects of the sodium ion.

**Figure 5.**
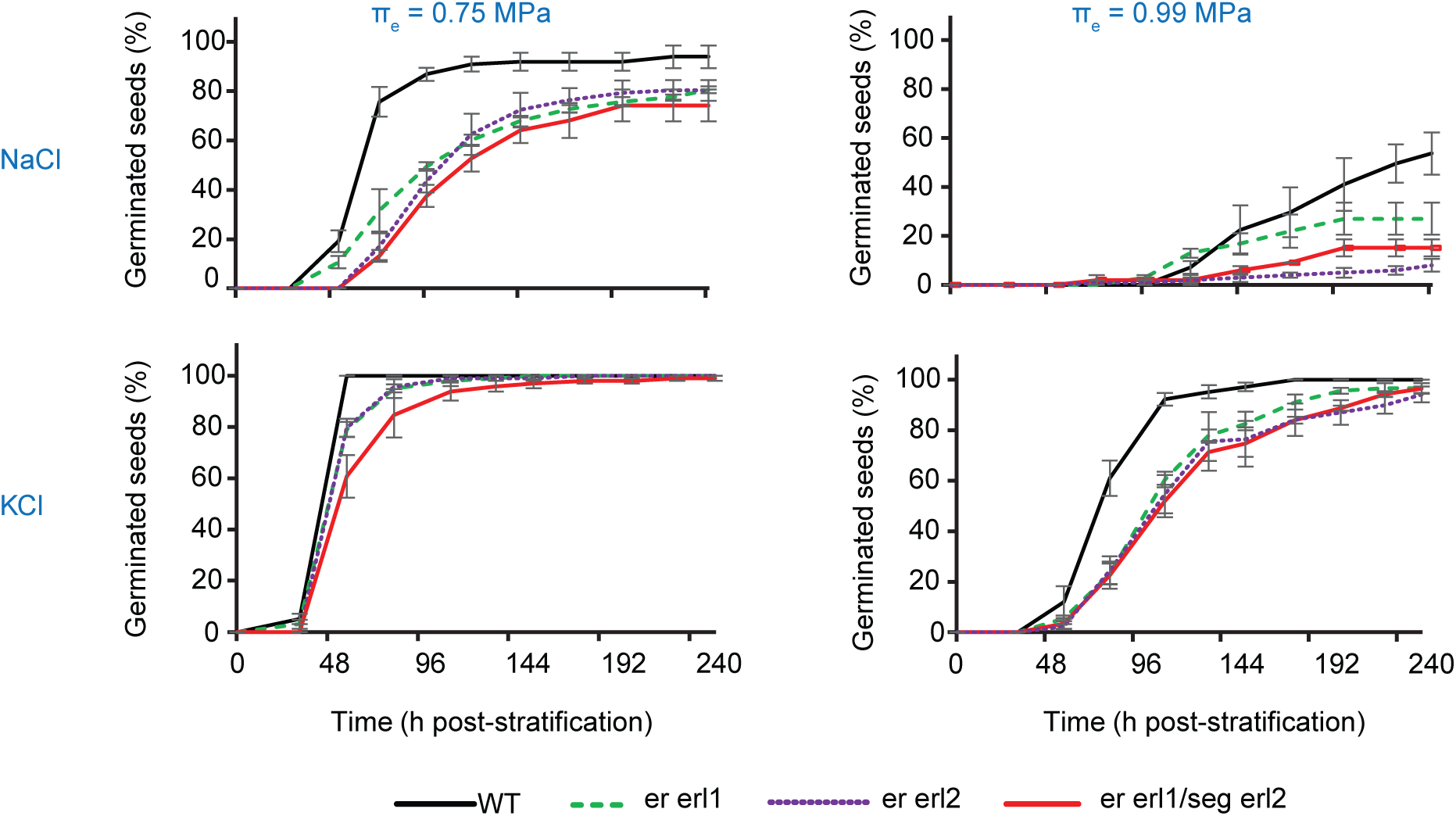
Seed germination in the salinity hypersensitive *er, er erl1, er erl2* and *er erl1/seg erl2* mutants is more sensitive to external NaCl than KCl concentrations under iso-osmotic conditions. Time-course of germination under iso-osmotic conditions (media osmotic pressure, πe) induced by supplementation of either NaCl or KCl. Percentages of seeds exhibiting endosperm rupture 4 d post-stratification (means and s.e.m.; n = 3 plates, 30 seeds per plate and per genotype; experiments replicated twice).

Notably, while all or the vast majority of WT, *erl1, erl2, erl1 erl2* seeds plated on NaCl medium eventually germinated (90 to 100%, similar to salt-free media), a significant proportion of *er, er erl1, er erl2* and *er erl1/seg erl2* seeds failed to do so, even after a lengthy incubation period (Fig. 6; Supplementary Fig. S1). Among those, a majority (up to 70%) did not even exhibit testa rupture. To test whether these were damaged or dead seeds, we transferred them to NaCl-free media. Most germinated readily, within 20-25 h (Fig. 6), bringing the final percentage of germinated seeds to similar levels as those observed for seeds never exposed to salt. Failure to germinate on saline media was thus not due to irreversible cellular damage and loss of seed viability, but rather to a slower or halted progression of the germination process. Consistent with their maintained viability and fast germination upon salinity stress release, seeds with arrested germination on salty media showed similar *ER*f expression levels as germinated seeds (*ERL1* and *ERL2* genes) or even higher (*ER*), (Fig. 3, comparison of III-NG to III-G seeds).

**Figure 6.**
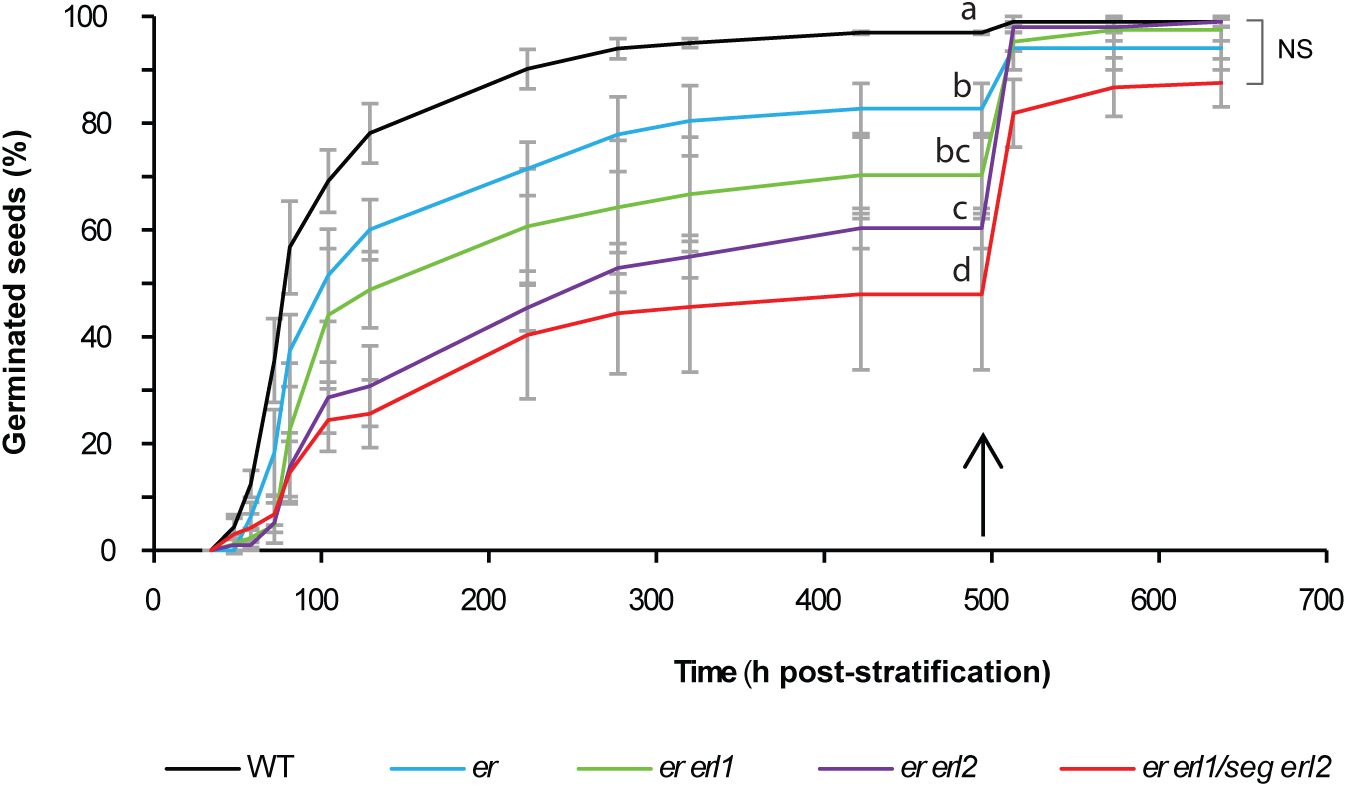
Seed germination readily resumes upon salinity stress removal. Time-course of seed germination on 150 mM NaCl media (0-490 h), and after transfer to NaCl-free media (arrow on x-axis). The experiment was repeated 3 times. Values are means and s.e.m. (n = 4 plates, 30 seeds per genotype per plate). Different letters above data points denote significant genetic differences at each time point by one-way ANOVA and Tukey HSD pair-wise tests (*P*□< □0.05). “NS” at the final time point indicates that genetic differences were non-statistically significant by one-way ANOVA.

### The ERf affects the ABA and GA regulation of seed germination

Salinity and osmotic stress promote ABA signalling and biosynthesis during germination (Seo et al., 2006; Piskurewicz et al., 2008; Yuan et al., 2010). ABA is a strong inhibitor of seed germination. To test whether the ERf-mediated differences in germination sensitivity to salinity stress are ABA-related, we compared germination kinetics of WT and *er*f seeds in the presence of ABA. ABA treatment consistently had a mild delaying effect on testa rupture, which was most pronounced for *er erl1* seeds (Fig. 7A). ABA strongly inhibited endosperm rupture also in an ERf-dependent manner (Fig. 7B). *er erl1* seed germination was the most sensitive to ABA, lagging behind WT even in the 1µM ABA range. Under higher ABA concentrations, seeds of the other two salt-hypersensitive mutants, *er erl2* and *er erl1/seg erl2*, but not *er*, also separated from WT, showing enhanced ABA sensitivity. Interestingly, so did the salinity non-hypersensitive *erl1 erl2* seeds (Fig. 7B). These data indicate the involvement of both ABA-dependent and ABA-independent pathways in the ERf-mediated sensitivity of seed germination to salinity.

**Figure 7.**
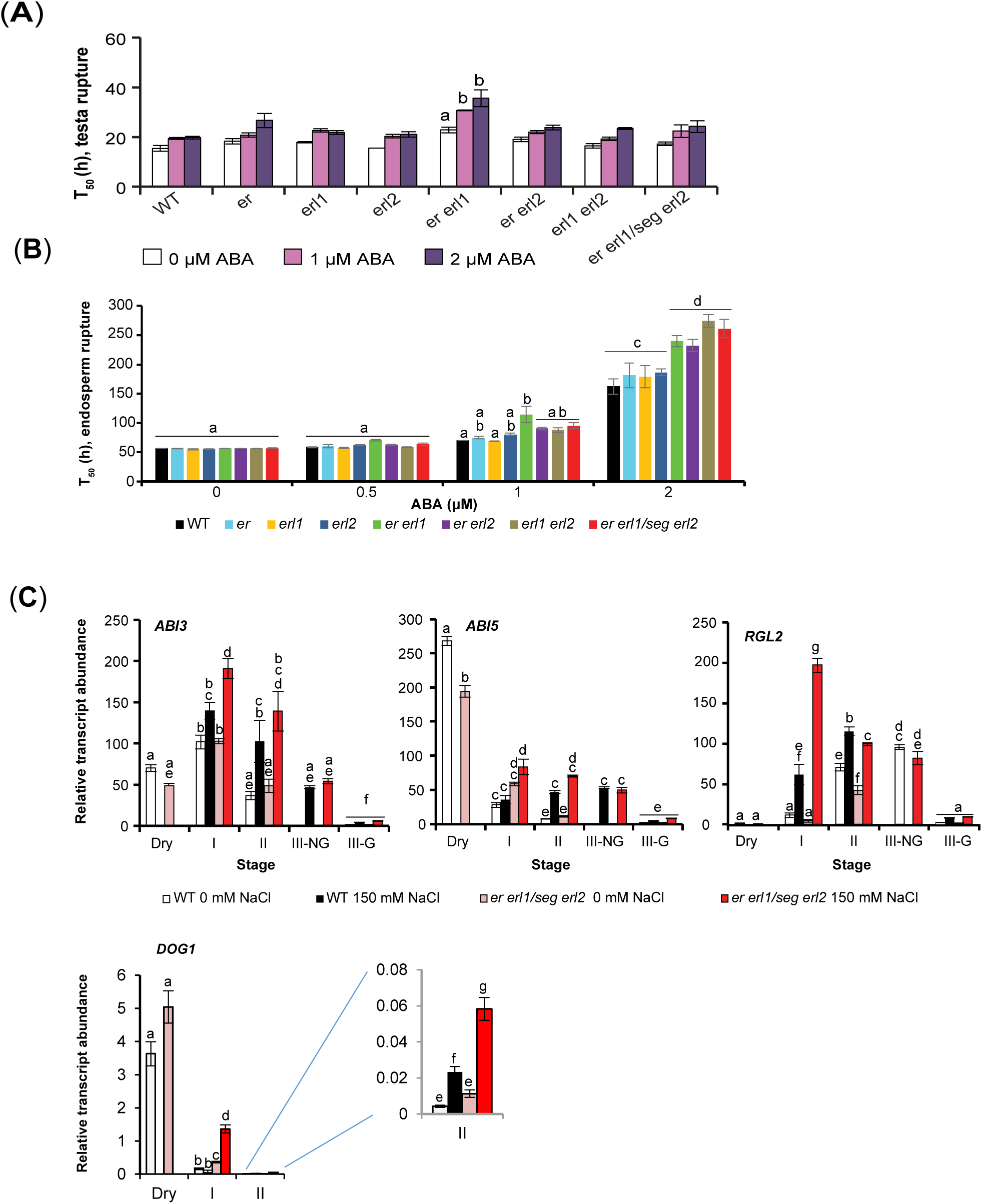
The ERf interacts with the sensitivity of seed germination to exogenous ABA and with the expression of major ABA and GA signalling genes. **A**, Germination response to exogenous ABA application. Data points represent T_50_ values and s.e.m. for testa rupture (**A**) and endosperm rupture (**B**) (n = 3 plates, with 30 seeds of each genotype). The experiment was repeated 3 times. **C**, *ABI3, ABI5, RGL2* and *DOG1* gene expression in dry seeds (“Dry”) and seeds sampled at: the end of imbibition and stratification (stage I); 20 h later (stage II, testa rupture); and 52 h later (stage III-G, endosperm rupture). Non-germinated seeds on 150 mM NaCl media (label III-NG) were analysed separately. **A**-**C**, Different letters indicate significant differences by two-way ANOVA and Tukey HSD pair-wise tests (*P*□< □0.05).

The germination inhibiting effect of ABA is antagonised by GAs (Koornneef et al., 1982; Holdsworth et al., 2008; Weitbrecht et al., 2011; Liu et al., 2016). Rather than the absolute levels of these hormones, the ABA/GA balance is key to the commitment of seeds to germinate. The DELLA RGL2 protein plays a pivotal role in the cross-talk between ABA and GA signalling in the imbibed seed. RGL2 acts as the main GA signalling repressor through activation of a number of transcriptional regulators, including ABI3 and ABI5, the central effectors of ABA signalling, establishment of dormancy, and repression of seed germination (Lopez-Molina et al. 2001, 2002; Lee et al., 2002; Piskurewicz et al. 2008, 2009; Liu et al. 2016). ABI3 and ABI5 are also involved in the regulation of early seedling growth arrest under water stress in Arabidopsis (Lopez-Molina et al. 2001; 2002), and in the reversible inhibition of germination in related *E.salsugineum* under salinity (Kazachkova et al., 2016). To better understand the interaction of the ERf with the ABA regulation of seed germination we monitored the expression of *ABI3, ABI5* and *RGL2* in WT and *er erl1/seg erl2* seeds during germination, and also of *DELAY OF GERMINATION1* (*DOG1*), a pivotal seed dormancy gene which genetically interacts with ABI3 and with a central type 2C protein phosphatase of the ABA signalling pathway during germination, and also regulates ABI5 expression (Dekkers et al. 2016; Née et al., 2017; Nishimura et al. 2018). Constitutive gene expression levels were similar in WT and mutant seeds. Salinity systematically caused an up-regulation of gene expression, but that was stronger in *er erl1/seg erl2* seeds than WT (Fig. 7C). This result indicates that the ERf-mediated signalling cascade of salinity interacts with the ABA-GA signalling network of germination and dormancy. We also examined the expression of ABA and GA biosynthetic genes - *ABA2* and *NCDE4*; *GA3OX1* and *GAOX2-*respectively. None showed a differential response to salinity between mutant and WT (Supplementary Fig. S5).

### The role of the ERf in seed germination partly overlaps with a role in seed size and is primarily maternally controlled

Seed germination occurs when the pressure exerted by the turgid expanding embryo radicle overcomes the mechanical resistance of the surrounding testa and micropylar endosperm (Linkies et al., 2009; Nonogaki, 2014). As *er erl1 erl2* mature embryos have smaller cotyledons (Uchida et al., 2013), we reasoned that reduced growth potential could be a factor in the delayed radicle emergence observed in that mutant and possibly the other salt-hypersensitive *er, er erl1, er erl2* mutants under salinity and osmotic stress. As a first step to examine this, we measured seed size as a surrogate for embryo size, since the *Arabidopsis* embryo occupies most of the seed volume. *er, er erl1, er erl2* and *er erl1/seg erl2* seeds, i.e. all salt-hypersensitive seeds, were significantly smaller than WT or *erl1* and *erl2* seeds, and even smaller than *erl1 erl2* seeds which were larger than WT (Fig. 8A). These data uncover a function of the ERf in seed size determination. They also suggest a link between the ERf function in germination sensitivity to salinity and its influence on seed size. However, the fact that *erl1 erl2* seeds germinate simultaneously with WT seeds in the presence or absence of salt despite their significant seed size difference, indicates that the link is not absolute.

**Figure 8.**
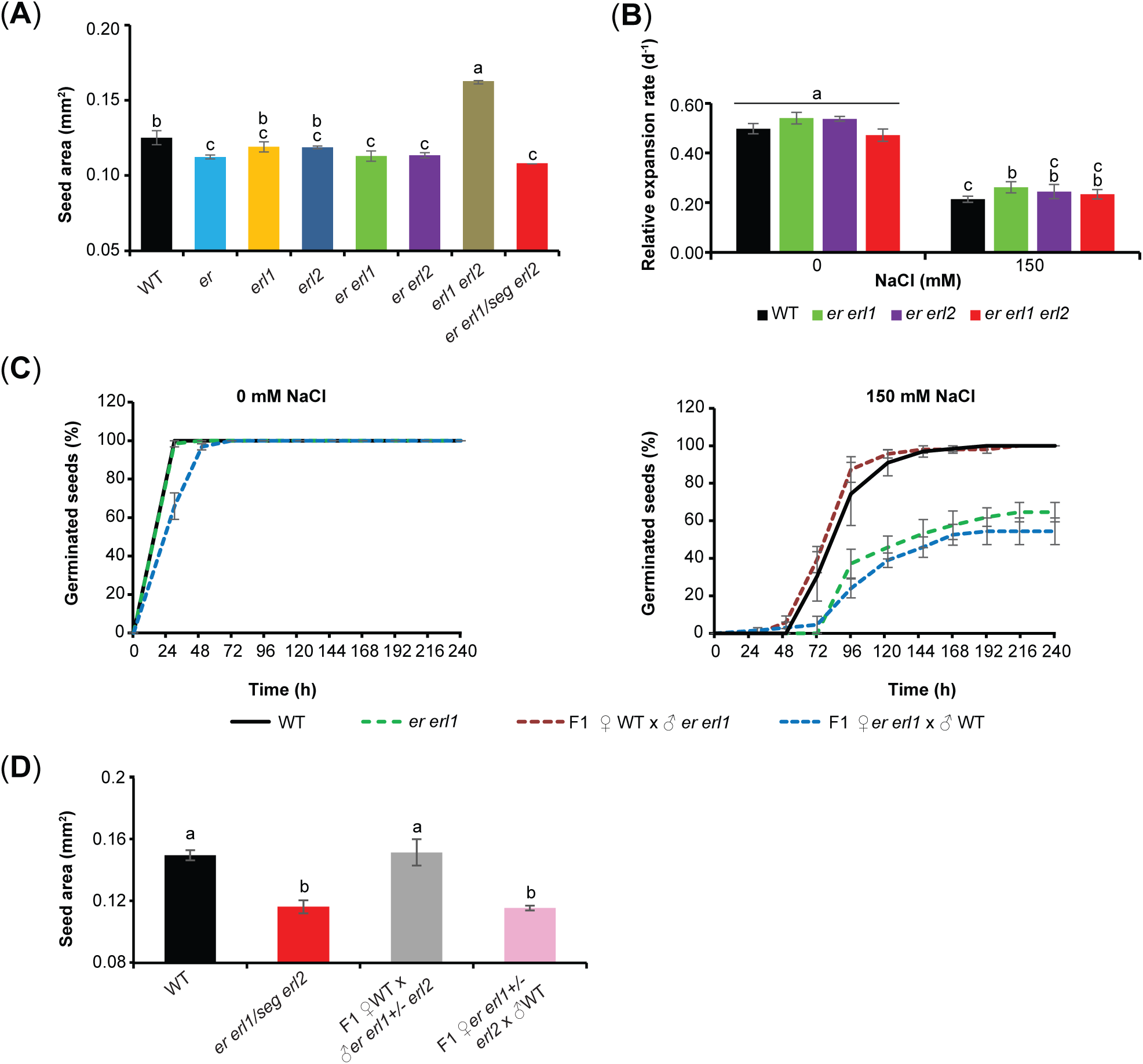
The ERf function in seed germination sensitivity to salinity is maternally controlled and shows partial overlap with an ERf function in the determination of seed size. **A**, Seed projected area (mm^2^); means and s.e.m. (n≥ 400 seeds per genotype, from 11 siliques). Letters indicate significant differences by one-way ANOVA and Tukey HSD pair-wise tests (*P*□< □0.001). **B**, Relative expansion rate (mm^2^ mm^-2^ d^-1^) of mature embryos excised from enclosing tissues, over a 72 h incubation period on 0 or 150 mM NaCl media, (n= 7). Different letters indicate significant differences by two-way ANOVA and Tukey HSD pair-wise tests (*P*□< □0.001). **C**, Time-course of germination for WT and *er erl1* seeds, and F1 seeds generated from their reciprocal crosses. Similar results were obtained from crosses between WT and *er erl2* flowers (data not shown). **D**, Size of F1 seeds from reciprocal crosses between WT and *er erl1+/-erl2* flowers (n=86 to 143 seeds per cross). Different letters indicate significant differences by one-way ANOVA and Tukey HSD pair-wise tests (*P*□< □0.001). **C-D**, Crosses were made between flowers at similar positions on the main inflorescence; seeds were harvested at the same time, 3 weeks after crossing.

We next considered the possibility of developmental defects in the smaller, salinity hypersensitive *er*f seeds. As expected, homozygous *er erl1 erl2* segregants displayed reduced, rounder cotyledons and a broader shoot apical meristem, as previously reported (Uchida et al. 2013). However, their hypocotyl and embryonic root were similar to WT, in length, number and size of constitutive cells (Supplementary Fig. S6).

Seeds reserves are essential for successful germination, and in Arabidopsis are mostly stored in cotyledons. Smaller seeds and cotyledons suggest less reserves, which could be responsible for hypersensitivity to salinity and osmotic stress. To examine this, we quantified fatty acid methyl esters (FAMES) derived from embryo lipids, which constitute the major fraction of Arabidopsis seed reserves (Penfield et al., 2004; Lionen and Schwender, 2009). There was no significant genetic difference across the range of genotypes, except for *er erl1/seg erl2* seeds (15% decrease) and thus, apart from that genotype, no correlation with germination sensitivity to salt (Supplementary Fig. S7A). The relative proportions of FAMES species were also similar across genotypes (Supplementary Fig. S7B-C). Taken together, these results indicate that the delayed or arrested germination of *er, er erl1, er erl2* and *er erl1/seg erl2* seeds on saline media was not likely due to reduced embryo size and growth potential *per se*.

Germination involves complex communication between embryo, seed coat, and intermediate endosperm –a one cell thin layer in the Arabidopsis seed. We therefore next considered a role of the ERf on seed germination via effects on the embryo surrounding tissues. To investigate that, we took advantage of the different contributions of the maternal and paternal genomes to the genetic make-ups of the three seed compartments (seed coat ♀ ♀, endosperm ♀ ♀ ♂, embryo ♀ ♂) and performed reciprocal crosses between WT and the salt-hypersensitive *er erl1* or *er erl2* mutants. These generated F1 seeds with same embryo genotype, but either WT or mutant seed coat, and predominantly WT or mutant endosperm. The two groups of F1 seeds germinated synchronously on NaCl-free media, but according to significantly different kinetics when challenged with salinity stress (Fig. 8C). Remarkably, for each cross, F1 seed germination occurred synchronously with seeds of the maternal parent. This result demonstrates that the function of the ERf in the regulation of germination sensitivity to salinity is primarily maternally controlled and mediated by the embryo-surrounding tissues, in particular the seed coat. Supporting this, when excised from their covering layers, “naked” *er erl1, er erl2* and *er erl1 erl2* mature embryos grew at similar rates as WT embryos, whether cultured with or without salt (Fig. 8B). F1 seeds also clustered with their maternal parent on seed size (Fig. 8D), showing the ERf effect on seed size is of maternal origin too, and strengthening the case for overlap of the ERf-dependent controls of seed size and germination response to salinity.

### The ERf-mediated regulation of seed germination involves the seed coat mucilage

Considering what properties of the seed coat the ERf might control to influence germination in a salinity-dependent manner we first tested for a role in seed coat permeability. To that end, seeds were incubated in tetrazolium red, a cationic dye classically used to detect seed coat defects and abnormal permeability (Wharton, 1955; Molina et al., 2008). Similar staining and tetrazolium salt reduction rates were observed across lines, except for significant increases in *er erl1* and to a small extent in *erl1* seeds (Fig. 9A), suggestive of increased seed coat permeability or NADPH-dependent reductase activity in these two mutants. We thus next measured seed sodium contents after 24 h stratification with or without salt. They showed no significant genetic variation (Fig. 9B). These results indicate that the observed differential germination response to salt among *er*f seeds cannot be ascribed to differences in seed coat permeability and accumulation of sodium ions *per se*.

**Figure 9.**
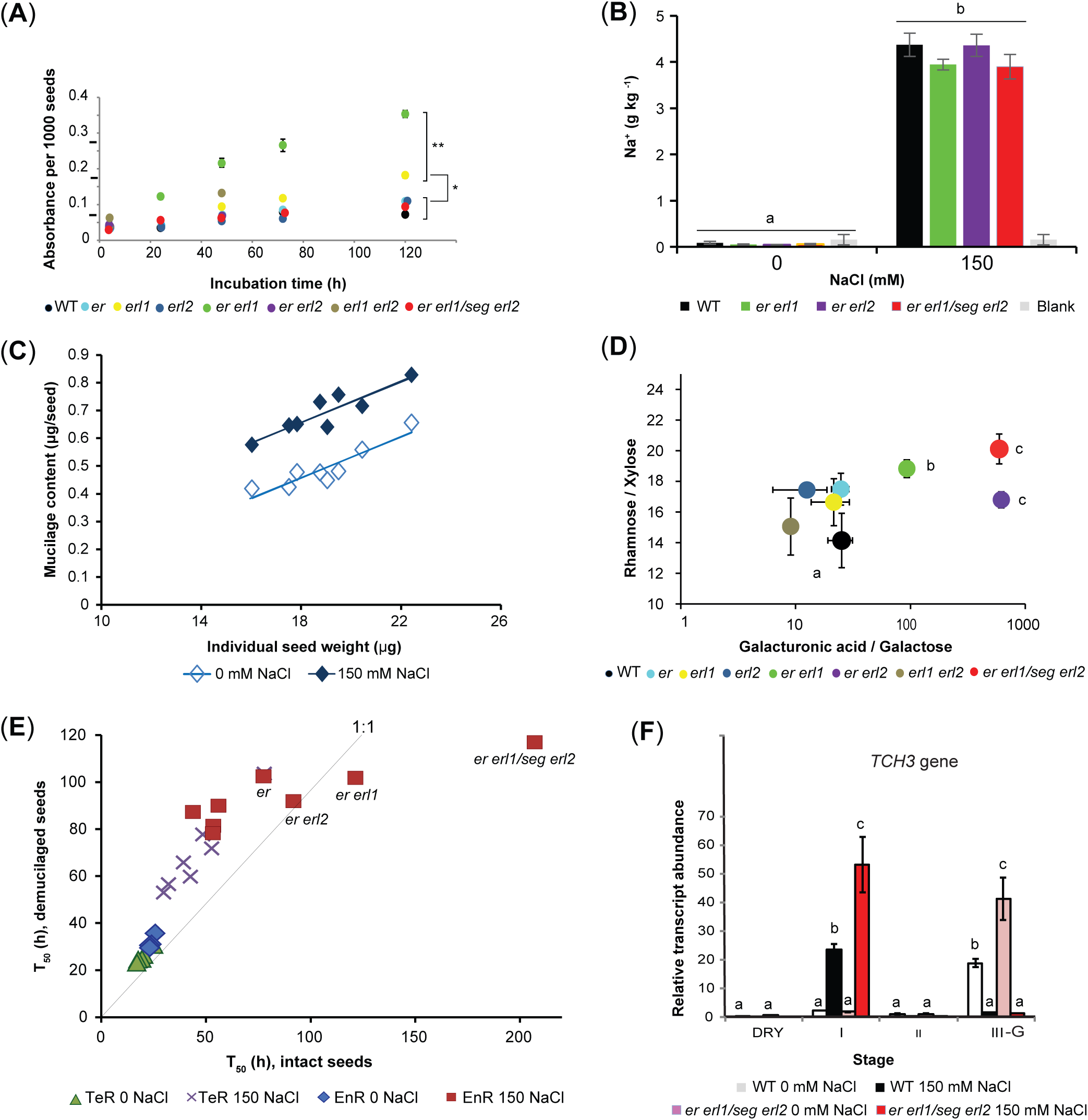
The ERf is involved in the control of seed coat permeability mucilage composition and salinity-dependent role in the regulation of germination speed. **A**, Seed coat permeability to tetrazolium red (n = 4 seed pools; some s.e.m. are hidden by symbols). * denotes statistical significance (P□<□0.001) by two-way ANOVA and Scheffe post-hoc test. **B**, Seed sodium content 24 h post-stratification on 0 mM or 150 mM NaCl media, (n=3 seed pools). Letters indicate significant differences by two-way ANOVA and Tukey HSD pair-wise tests (*P*□= 0.42 and 0.39 for genotype effect under control and salt treatment, respectively). **C**, Correlation between mass of water soluble mucilage per seed and seed size. Means and s.e.m. (n = 4 seed pools per genotype, 40 mg seeds per pool, average seed weight and area determined on sub-aliquots; experiment replicated 3 times). Regression lines: 0 mM NaCl, y=36.6x-0.20, r^2^ =0.84; 150 mM NaCl, y=36.3x+0.002, r^2^ =0.81. Similar results were obtained with size expressed as area**. D**, GalUA/Gal and Rhm/Xyl ratios. Letters besides points indicate statistical significance of differences in GalUA/Gal (*P*<0.05) by one-way ANOVA and Tukey post-hoc tests, compared to all unlabelled data points. *P*=0.08 for differences in Rhm/Xyl between *er erl1/seg erl2* and WT. **E**, Testa rupture (TeR) and endosperm rupture (EnR) T_50_ values for intact seeds and “demucilaged” seeds. Mean values per genotype (n = 3 plates; 30 seeds per genotype per plate). Labelled points denote genotypes where removal of the outer water soluble mucilage significantly advanced germination on 150 mM NaCl media. The 1:1 line represents the bisextrix, where mucilage removal is neutral. **F**, *TCH3* gene expression in WT and *er erl1/seg erl2* dry and imbibed seeds during the three germination phases, n=4 seed pools per genotype and NaCl condition, of 300 seeds each.

During seed coat differentiation on the mother plant, the specialised epidermal cells secrete mucilage polysaccharides that line their inner walls and build a central volcano-shaped columella (Beeckman et al., 2000; Western et al., 2000; Haughn and Western, 2012). Upon hydration, the desiccated, highly hydrophilic mucilage rapidly swells and ruptures the enclosing outer primary wall, wrapping the seed in a gelatinous capsule traversed by cellulosic rays radiating from the columella. Mutant seeds affected in mucilage synthesis or extrusion have been reported to be more sensitive to low water potential during germination (Penfield et al., 2001; Yang et al., 2010). This prompted us to next examine mucilage release by WT and *er*f seeds upon imbibition. We collected the loosely adhering mucilage which can easily be detached from the seed surface, as opposed to the inner, cell wall-bound fraction. Large genetic variation was observed in the amounts recovered, but that scaled with genetic variation in seed size (Fig. 9C). Salinity caused a large increase in mucilage extrusion, but of similar proportion in all genotypes, resulting in a simple translation of the relationship observed on control media. Consistently, mucilage staining with ruthenium red, a classic dye that binds pectins, showed a thicker and often darker mucilage halo under saline than control conditions, but with no indication of variation among genotypes within each treatment (Supplementary Fig. S8A, B).

Recent studies suggest the importance for germination of the mucilage physico-chemical properties and attachment to the seed rather than simply its amount (Rautengarten et al., 2008; Saez-Aguayo et al., 2013). We thus next analysed mucilage composition. The expected sugars were detected, mostly rhamnose and galacturonic acid (GalUA) derived from rhamnogalacturonans type I (RG I) - the major pectin of the *Arabidopsis* seed mucilage (Macquet et al., 2007; Arsovsky, 2009) - and low amounts of other neutral and acidic sugars, derived from RGI-s side chains (Supplementary Fig. 8C). When analysed individually, these sugars showed no statistically significant variations among genotypes. However, examination of compositional profiles by multivariate analysis suggested genetic variation in the relative abundance of backbone sugars (Rhm and GalUA) and some side-chain sugars (Xyl and Gal), leading us to compare their ratios across the full spectrum of lines (Fig. 9D). This revealed dramatically increased GalUA/Gal ratios in *er erl1, er erl2* and *er erl1/seg erl2* mucilage compared to WT and other lines (*P*=0.027), and a trend for higher rhamnose to xylose ratios in mutant mucilage other than *erl1 erl2*, especially in *er erl1* and *er erl1/seg erl2* mucilage (*P*=0.08). These results suggest that the ERf plays a role in the control of mucilage composition and architecture *via* interactions with the mechanisms controlling the abundance of carboxyl sites - i.e. of potential sites for pectin cross-linking - and perhaps also pectin branching. Moreover, they indicate a link between such a role and the ERf function in the regulation of seed germination.

To test this and probe causality, we took an indirect, holistic approach, and compared the germination kinetics of intact seeds and demucilaged seeds, deprived of the shell of loosely adherent mucilage extruded during imbibition. Demucilaged seeds systematically germinated more slowly than intact seeds on salt-free media (Fig. 9E), as is common. Under saline conditions, this was also the case for WT, *erl1, erl2* and *erl1 erl2* seeds but, strikingly, in *er erl1, er erl*2 and *er erl1/seg erl2* seeds, mucilage removal had the opposite effect: germination delay behind WT seeds was reduced, with radicle protrusion lagging by 23 h, 13 h and 39 h behind WT, respectively, instead of 68 h, 39 h, and 153 h, respectively, for intact seeds (Fig. 9E). This reflected faster progression from testa rupture to endosperm rupture. These data demonstrate a critical role of the seed water soluble mucilage in mediating the salinity-dependent function of the ERf in controlling the completion of seed germination.

Although appearing as distinct layers upon imbibition, mucilage and cell walls are tightly bound. The suberised seed coat and underlying endosperm constitute a mechanically strong barrier that needs to be weakened to enable radicle emergence. The micropylar endosperm that surrounds the radicle tip is thought to be the major source of mechanical resistance to radicle protrusion (Linkies et al., 2009; Dekkers et al., 2013). Endosperm weakening is effected by cell wall-modifying enzymes, in interaction with ROS and hormonal signals from the embryo, GA especially (Finch-Savage and Leubner-Metzger 2006; Muller et al., 2006; Penfield et al., 2006). We thus hypothesised that the importance of the mucilage and seed coat in mediating delayed or arrested germination in the *er, er erl1, er erl2* and *er el1/seg erl2* mutants on saline media, could in part be related to ERf-dependent differences in endosperm and seed coat mechanical properties. The *Arabidopsis* seed is too small for direct measurements of testa and endosperm rupture forces, as is possible in other species (Linkies et al., 2009), leading us to instead examine the expression of the *Arabidopsis TOUCH (TCH*) gene *TCH3*, which encodes a calmodulin-like protein and is greatly up-regulated in response to a range of mechanical signals in other tissues (Braam and Davis, 1990). Comparison of *TCH3* expression in WT and *er erl1/seg erl2* seeds (Fig. 9F) showed the presence of transcripts in dry seeds, at similar, low levels. Imbibition triggered de novo *TCH3* transcription on 150 mM NaCl media in both WT and *er*f mutant seeds, consistent with the known role of calcium in salinity signalling (Munns and Tester, 2008). Remarkably, that induction was significantly enhanced in *er erl1/seg erl2* seeds, and was transient, preceding testa and then disappearing. These data are suggestive of enhanced mechanical constraint imposed on *er erl1/seg erl2* than WT embryos before endosperm rupture. On control media, de novo *TCH3* transcription did not occur before the final phase of germination phase; again it was enhanced in *er erl1/seg erl2* seeds compared to WT.

## Discussion

Plant propagation, dispersion, ability to compete and yield all ultimately rely on viable seeds being produced and able to germinate at a time favourable to autotrophic growth and establishment of a new seedling. During development on the mother plant, following embryogenesis and acquisition of dormancy during maturation, the seed undergoes intense dehydration and the embryo becomes quiescent. Germination brings that embryo from a highly resilient to a highly vulnerable state, in direct contact with the outer environment, and to a point of no return. How seeds monitor conditions in their immediate surrounding to optimise the timing of germination initiation and its completion is mostly unknown. In this study, we show that the *Arabidopsis* ERECTA family acts to control the timing of seed germination according to external salinity and osmotic levels (Fig. 1; Fig. 4). Loss of *ER*, or of *ER* and its paralogs slows down germination or even prevents it under increasing salinity and osmotic stress, while not compromising seed viability, as germination readily resumes upon the return of favourable conditions (Fig. 6). The ERf-mediated sensing of changing salinity levels involves interactions with the ABA-GA signalling network of germination and dormancy, and is primarily controlled by the embryo surrounding endosperm and testa, with a critical role of the latter and its mucilage (Fig. 8; Fig. 9). These findings reveal unsuspected regulators of the interactions between the seed and its environment, and a novel function of the three ERf receptor-like kinases in controlling these interactions, and cryptic genetic variation in seed germination.

### The *ERECTA* gene family regulates seed germination on salt, via maternally controlled effects on seed coat enlargement and mucilage properties

The seed coat derives from the maternal ovule integuments which, following fertilisation, expand and undergo profound developmental and biochemical transformations, resulting in a highly differentiated, impermeable and mechanically strong tissue (Beeckmann et al. 2000; Western et al. 2000). The mucilage is secreted and deposited in its outer, epidermal layer, concomitantly with embryo morphogenesis, following cessation of integument expansion (reviewed by Haughn et al., 2012; North et al., 2014). Its physiological roles have remained elusive. Apart from anchoring the imbibed seed to its physical substrate, the gelatinous mucilage is generally thought to facilitate germination, especially under osmotic stress, through sequestering water and keeping the seed hydrated (Penfield et al., 2001; Arsovski et al., 2010; Yang et al., 2010). However, several studies suggest mucilage can also inhibit germination under unsuitable conditions, perhaps through limiting water and oxygen diffusion to the embryo (Western et al., 2012 and references herein). This study sheds some light on the ill-understood genetic control of the context-dependent role of the seed mucilage on germination, revealing that the ERf are key players. We observed a promoting role of the seed mucilage on germination speed in WT, *erl1, erl2* and *erl1 erl2* seeds, under both saline and non-saline conditions (Fig. 9E). However, in the salt-hypersensitive *er erl1, er erl2*, and *er erl1/seg erl2* seeds, that role was only expressed under control conditions. Under salinity, it was lost (*er erl1, er erl2* seeds) or even reversed (triple mutant). These results link, for the first time, the seed mucilage and the *ER* pathway in the germination response to environmental variations at the seed surface.

How could the ERf control salinity-dependent properties of the seed mucilage regulating the germination process? The seed mucilage is alike a pectin-rich secondary cell wall (Haughn and Western, 2012). The degrees of pectin branching and cross-linking, with calcium ions in particular, are known to greatly influence pectins’ hydrophilicity, adsorption to cellulose microfibrils and partitioning between loose outer mucilage and adherent inner mucilage (Willats et al., 2006; North et al., 2014; Ralet et al., 2016). It is also well-established that the small monovalent Na^+^ ions have the capacity to easily displace the larger divalent Ca^2+^ ions that cross-link carboxyl residues of adjacent pectin molecules (Fry, 1986; Willats et al., 2006; Ghanem et al., 2010), thus leading to a looser, more hydrophilic mucilage upon imbibition with saline than salt-free water, and also more abundant (Fig. 9C; Ghanem et al. 2010) due to increased release of pectin molecules from the cellular matrix. We propose that the enrichment of *er erl1, er erl2*, and *er erl1/seg erl2* seed mucilage in uronic acids - hence potential sites for Ca^+2^ _-_ Na^+^ exchange- and trend to reduced xylose content relative to backbone rhamnose suggestive of altered branching (Fig.9D), thus have the potential to significantly modify a) the seed mucilage and sub-tending wall swelling properties and changes in osmotic potential, conformation and rigidity upon imbibition with a saline or high osmolarity solution (Willats et al., 2006; Ghanem et al., 2010; Ralet et al., 2016); b) the rearrangement of mucilage and wall components as pectin molecules get released (Rautengarten et al., 2008); and c) perhaps also free Ca^2+^ influx to the adjoining inner endosperm and embryo; so d) as a whole, the chemical and mechanical interactions between the seed environment, seed coat and interior compartments.

The *er*f seeds with enhanced sensitivity to salt and hyperosmotic stress during germination are also smaller (Fig. 8A). *Arabidopsis* seed size is controlled by complex interactions of zygotic and maternal factors, and seed integuments-endosperm inter-signalling (Garcia et al., 2003; Luo et al., 2005; Day et al., 2008; Dilkes et al., 2008; Zhou et al., 2009; Wang et al., 2010; Jiang et al., 2013). Here, our reciprocal crosses show that variation in final seed size among *er*f mutants and WT is of maternal origin (Fig. 8D). Final seed size is reached early in seed development, through a first phase of active cell proliferation triggered by fertilisation, in both the integuments and the endosperm, followed by a period of mostly cell expansion. Expansion ceases five to six days post-anthesis, concomitantly with the endosperm switching from syncitial development to cellularisation (Garcia et al. 2005), and the start of starch and mucilage synthesis. Variation in maximum cell elongation appears to be the main driver of maternal variations in final seed cavity and seed size as observed here, through a so-called ‘compensatory” growth mechanism (Garcia 2005). The *ERECTA* gene has been implicated in such compensatory mechanism between cell number and cell size in leaves (Ferjani et al. 2007), and comparison of the seed epidermis of the *Ler* and *Columbia* accessions suggests “compensation” may take place in the seed integuments too (Garcia et al. 2005). Interestingly, the progression and completion of integument growth during ovule development was previously reported to require a minimum ERf signalling (Pillitteri et al., 2007). That requirement was ascribed to a role of the ERf in cellular proliferative activity through interactions with cell cycle regulators. However, the final cell number in the mature ovule was unchanged making it unlikely that the reduced seed size cavity and less expanded seed coat observed here in the *er, er erl1, er erl2* and *er erl1/seg erl2* seeds (Fig. 8A) are pre-determined prior to fertilisation. Moreover, no ovule integument growth defect was reported in ovule integuments other than *er erl1 erl2+/-*. Here, loss of *ER* alone was sufficient to cause reduced seed size, and further loss of *ERL1* or *ERL2* had only a small or no significant additional inhibitory effect; and, when occurring in an *ER* background, loss of *ERL1* and *ERL2* instead caused an increase of seed size beyond that in WT (Fig. 8A). This supports the idea that partly different mechanisms are involve in ERf-mediated control of seed size and germination sensitivity to salinity. Given the role of the ERf in the composition of the mucilage (Fig. 9D) and the reported increases in uronic acids and cellulose in leaves of two *er* mutants (Sánchez-Rodriguez et al., 2009), a tentalising hypothesis is that the ERf may regulate cell wall formation and assembly, not only during mucilage and secondary cell wall deposition, but also prior to that, during seed coat enlargement and formation of the seed cavity. That proposition would provide a unifying explanation for a link between the ERf-mediated regulation of seed size, salinity-dependent mucilage properties and germination speed, as uncovered by this study.

### ERf-mediated salt signalling in germinating seeds involves a complex regulatory network

The *Arabidopsis* seed coat is in immediate contact with the one-cell thin endosperm, itself in direct contact with the embryo. Although less well-documented than in humans, there are demonstrated cases of plant membrane receptors or mechano-sensitive channels’ ability to monitor cell wall integrity, membrane and wall physical interactions, deformation and rheology (Hamann et al., 2012; Monshausen and Haswell, 2013; Hamilton et al., 2015; Haswell and Verslues, 2015). The ERf proteins belong to XIII Leucine-Rich Repeats Receptor-Kinases. Most interestingly, among its other four members, that class includes FEI1 and FEI2 (Shiu et al., 2004) which were recently shown to interact with an arabinogalactan protein in mediating a salt-overly sensitive root and seed adherence mucilage phenotype (Harpaz-Saad et al., 2011; Griffiths et al., 2014). In addition, based on the analysis of disease resistance in two *er* mutants, the ER protein has been suspected of interacting with wall-associated kinases (WAKs) during defence against some pathogens, via effects on cell wall composition (Sánchez-Rodríguez et al., 2009). WAKs are known to be tightly bound to pectins, galacturonic acids especially, in a Ca^2+^-dependent manner (Wagner and Kohorn, 2001; Decreux and Messiaen, 2005), and several WAK/WAK-Like proteins have been implicated in responses to mineral ions, including Na^+^ (Sivaguru et al., 2003; Hou et al., 2005; de Lorenzo et al., 2009), and to osmotic stress (SOS6/AtCSLD5, Zhu et al., 2010), through unknown mechanisms. ERf-mediated modifications of mucilage and bound cell walls may thus be perceived and signalled to the seed interior by the *ER*f proteins themselves either directly or through modified interactions with cell wall-associated proteins, osmo-sensors or mechano-sensors (Dekkers, et al. 2013; Nonogaki, 2014). The induction of *TCH3* in the *er erl1/seg erl2* seed supports this hypothesis.

It will be intriguing to unravel the downstream cascade. The salt-hypersensitive *er erl1, er erl2, er erl1/seg erl2* seeds show enhanced sensitivity to exogenous ABA, and enhanced upregulation *of ABI3, ABI5* and *RGL2* under saline conditions compared to wild type (Fig. 7). *ABI3, ABI5, RGL2* are emerging as important mediators of salinity and osmotic stress and controllers of ABA-GA homeostasis in imbibed seeds. ABA synthesised in the endosperm and released to the embryo activates the abundance and activity of the *ABI3* and *ABI5* transcription factors, and triggers an auto-feedback loop that maintains *RGL2* mRNA levels high and represses cell wall modifying enzymes (Giraudat et al., 1992; Finkelstein and Lynch, 2000; Lee et al., 2002; Lopez-Molina et al., 2001 & 2002; Piskurewicz et al., 2008; Piskurewicz et al., 2009; Lee et al. 2010; Kang et al. 2015). Our data indicate that the ERf-mediated regulation of germination sensitivity to changing salinity levels interferes with that signalling loop.

A well-documented adaptive mechanism seeds have evolved to withstand unfavourable conditions such as high temperatures, cold, osmotic or salinity stress, and maintain embryo viability is secondary dormancy (Bewley, 1997) - a reversible, transient quiescent state induced and released in adaptation to fluctuating environmental conditions (Koornneef et al., 1982; Giraudat et al., 1992; Léon-Kloosterziel et al., 1996; Finch-Savage and Leubner-Metzger, 2006; Lefebvre et al., 2006; Weitbrecht et al., 2011; Ibarra et al., 2016). ABI3, ABI5 and RGL2 are prominent players in the regulation of secondary dormancy and increased sensitivity to ABA, upregulation of *ABI3, ABI5* and *RGL2* have been reported during early growth arrest in newly germinated *Arabidopsi*s seedlings under water stress and salinity (Lopez-Molina et al., 2001; 2002). Here we find that loss of the ERf sensitises seed germination to salinity and frequently arrests it, and that this arrest is reversible, with germination readily resuming upon stress release and progressing to completion as fast as in seeds never exposed to stress (Fig. 6). Moreover, arrested seeds show an upregulation of the DOG1 gene (Fig. 7), a major controller of coat- and endosperm-mediated dormancy as takes place in the Arabidopsis seed. *DOG1* interacts with GA and ABA signalling, upstream of ABI5, and appears to be an agent of environmental adaptation of germination among *Arabidopsis* accessions (Graeber, 2014; Dekkers et al., 2013; Née et al., 2017; Nishimura et al. 2018). Taken as a whole, these observations suggest that the ERf interacts with the molecular controls of secondary dormancy to appropriately cue and pace germination. While promotion of fast germination under stress may be seen as desirable, it also exposes the newly germinated seedling to risks of death should adverse conditions persist or worsen as the embryo becomes directly exposed to the external environment with all its reserves already burnt. In such circumstances, germination delay or arrest could then be a useful protective strategy to maximise chances of survival through temporarily safeguarding the embryo against such a fate. In that light, the environment-dependent function of the ERf on germination speed would perform a vital adaptive function. Interestingly, only under extremely severe stress (∼200 mM NaCl) does the loss of *ER* and *ERL1* and/or *ERL2* cause germination arrest in absolutely all seeds within a cohort. Under milder stress, some seeds do germinate at the same time as WT, others with increasing delay, and others are arrested until stress release, a mixed response that may balance risks of death and loss of fitness or ability to complete the life cycle in time.

In conclusion, plants must be endowed with a “surveillance” system for the perception and transduction of external environmental cues to internal compartments, and their integration with developmental pathways. This study illuminates a key role of the ERf in that elusive integrative network in seeds, to control the most critical decision in the cycle of life, when to initiate a new plant. Given the evolutionary conservation of the ERECTA receptor-kinases across a broad range of plant species, and the emerging intense interest in mucilage as a model for cell wall studies and an important adaptive feature, our findings open new avenues for unravelling the mechanisms seeds have evolved to control germination and tune it to local conditions for maximising chances of survival.

### Gene accession numbers

*AtER (At2g26330), AtERl1 (At5g62230), AtERl2 (At5g07180)*, (*At4g37490*), *AtPDF2* (*At1g13320), bHLH* (At4g38070), *PPR* (At5g55840)

## Supplementary Data

**Supplementary Figure S1.** NaCl-dependent effects of reduced ERf signaling on the time course of seed germination.

**Supplementary Figure S2.** Loss of *ER* alone or in combination with *ERL1* and *ERL2* sensitises seed germination to salinity in a dose-dependent manner.

**Supplementary Figure S3.** ERf-dependent sensitivity of seed germination to NaCl in an independent set of *er*f single, double and triple knock-out mutants.

**Supplementary Figure S4.** *ER*f promoter activity in mature dry seeds and germinating seeds.

**Supplementary Figure S5.** Similar sensitivity to salinity stress of selected ABA and GA biosynthetic gene expression levels in WT and *er erl1/seg erl2* seeds.

**Supplementary Figure S6.** Mature *er erl1 erl2* embryos exhibit similar radicle size and patterning than WT seeds.

**Supplementary Figure S7.** Relative abundance of total fatty acid methyl-esters (FAMES) and relative proportions of individual species in embryos at full seed maturity.

**Supplementary Figure S8.** Characteristics of the seed mucilage.

**Supplementary Table S1.** List of genotyping and RT-qPCR primers.

## Acknowledgments

We thank Josephine Ginty and Kefan Peng for assistance with seed permeability assays and quantitative RT-PCR, respectively; Richard Phillips for help with measurements of seed ion contents; Guillaume Tcherkez for discussions on metabolites multivariate analysis; Keiko Torii for seeds of *proERf:GUS* reporter lines; the Nottingham Arabidopsis Stock Centre and SALK-Institute for mutant seeds; the Australian National University for funding.

---

**Supplementary Figure S1. NaCl-dependent effects of reduced ERf signaling on the timing and pace of seed germination.**

**A-D**, Percentages of seeds exhibiting testa rupture (**A, C**) and endosperm rupture (**B, D**) on 0 mM NaCl (**A, B**) and 150 mM NaCl media (**C, D**) as a function of time (h post-stratification). Data points are means and s.e.m. (n = 4 plates per condition, 30 seeds per genotype in each plate). The experiment was repeated 5 times.

**Supplementary Figure S2. Loss of *ER* alone or in combination with *ERL1* and *ERL2* sensitises seed germination to salinity in a dose-dependent manner.**

**A**, Percentages of seeds exhibiting endosperm rupture 4 d post-stratification on 0, 100, 150 or 200 mM NaCl (means and s.e.m.; n = 3 plates per condition, 30 seeds per genotype per plate). The experiment was repeated 3 times. **B**, Percentages of germinated seeds over a 10 d incubation period on 200 mM NaCl media for WT and the four NaCl-hypersensitive *er*f mutants. Different letters indicate significant differences by one-way ANOVA and Tukey HSD pair-wise tests (*P*□< □0.001), n = 4 plates).

**Supplementary Figure S3. ERf-dependent sensitivity of seed germination to NaCl in an independent set of *er*f single, double and triple knock-out mutants.**

**A**, *ERL1* and *ERL2* expression is abolished in the *erl1-5* (SALK_019567), and *erl2-2* (SALK_015275C) mutants. **B**, T_50_ values for testa and endosperm rupture in WT and *er*f mutants carrying the *er2, erl1-5*, or *erl2-2* alleles. **C**. Time-interval between testa and endosperm rupture. **A-C.** Means and s.e.m. are shown (n = 4 plates, 30 seeds per genotype in each plate; experiment replicated 3 times). Different letters indicate significant differences by two-way ANOVA and Tukey HSD pair-wise tests (*P*□< □0.001).

**Supplementary Figure S4. *ER*f promoter activity in mature dry seeds and germinating seeds. A-C**, GUS staining patterns for embryos dissected from mature dry seeds, just before sowing (**A**), and then from germinating seeds incubated on 0 mM NaCl media (**B**) or 150 mM NaCl media (**C**), at the end of stratification (0 h time point) and daily thereafter until all seeds had germinated.

**Supplementary Figure S5. Similar sensitivity to salinity stress of selected ABA and GA biosynthetic gene expression levels in WT and *er erl1/seg erl2* seeds.**

*ABA2, NCED9, GA3OX1* and *GA3OX2* relative gene expression in dry seeds (“Dry”), after imbibition and stratification (I), and at the end of the next two germination phases (stages II, III; see Methods; III-NG non-germinated seeds yet on 150 mM NaCl media). n=4 pools of seeds per genotype and media, of 300 seeds each). Different letters within each graph indicate significant differences by two-way ANOVA and Tukey HSD pair-wise tests (*P*<0.01).

**Supplementary Figure S6. Mature *er erl1 erl2* embryos exhibit similar radicle size and patterning than WT seeds.**

**A**, Representative photographs of WT and *er erl1 erl2* mature embryos excised from dry seeds dissected out of siliques of the same age and similar positions on the main inflorescence. Embryos were cleared and imaged by differential interference microscopy. **B-E**, Morphometric analysis: cotyledon and radicle lengths (**B**); number of hypocotyl cells (**C**); cotyledon size (**D**); and number of cotyledon epidermal cells (**E**). * denotes significant differences by 2-tailed paired t-tests (*P* = 0.0011; n=5).

**Supplementary Figure S7. Relative abundance of total fatty acid methyl-esters (FAMES) and relative proportions of individual species in embryos at full seed maturity. A**, Amount of FAMES in mature embryos. Data points show amounts for 4 independent pools of 50 embryos for each genotype. Genotypes sharing a letter above the box are not statistically different by one-way ANOVA and Tukey pair-wise tests (*P*□<□0.05), **B**, Percentages of medium-chain fatty acids (C16, C18; tall bars, dark shade colors) and mono-unsaturated fatty acids (shorter bars, paler shade colors); the complements to 100% represent long-chain fatty acids (C20,C22) and polyunsaturated fatty acids, respectively. **C**, Proportions of individual fatty acids relative to the total amount of FAMES. Measurements were done by GC-MS on 4 pools of 50 mature embryos excised from dry seeds.

**Supplementary Figure S8. Characteristics of the seed mucilage. A, B**, Ruthenium red staining of WT and *er*f seed mucilage after 2 h imbibition in 0 mM NaCl (top row) or 150 mM NaCl solution (bottom row), with gentle shaking (**A**) or in presence of 10 mM Tris-HCl without shaking (**B**). Ruthenium red stains acidic pectins. **C**, Proportions (%) of monosaccharides in seed water soluble mucilage (n = 4 pools of seeds, 40 mg seeds per pool). Genetic differences for individual sugars were not statistically different by one-way ANOVA and Tukey HSD pair-wise tests at *P*<0.05, but *P*=0.055 to 0.08 for differences in xylose content between WT and *er, erl1, er erl1, er erl2* and *er erl1/seg erl2* mucilage. Rhm, rhamnose; Ara, arabinose; Xyl, xylose; Man, mannose; Gal, galactose; GalUA, Glucoronic acid measured in the water soluble mucilage, which typically represent 60-70% of the total Arabidopsis seed mucilage (Ralet et al. 2016).

**Supplementary Table S1. List of genotyping and RT-qPCR primers**

